# Grafting or Pruning in the Animal Tree: Lateral Gene Transfer and Gene Loss?

**DOI:** 10.1101/229468

**Authors:** Julie C. Dunning Hotopp

## Abstract

Lateral gene transfer (LGT) into multicellular eukaryotes with differentiated tissues, particularly gonads, continues to be met with skepticism by many prominent evolutionary and genomic biologists. A detailed examination of 26 animal genomes identifed putative LGTs in invertebrate and vertebrate genomes, concluding that there are fewer predicted LGTs in vertebrates/chordates than invertebrates, but there is still evidence of LGT into chordates, including humans. More recently, a reanalysis a subset of these putative LGTs into vertebrates concluded that there is not horizontal gene transfer in the human genome. One of the genes in dispute is an N-acyl-aromatic-L-amino acid amidohydrolase (ENSG00000132744), which encodes ACY3, which was initially identified as a putative bacteria-chordate LGT but was later debunked has a significant BLAST match to a more recently deposited genome of *Saccoglossus kowalevskii,* a flatworm, Metazoan, and hemichordate. Using BLAST searches, HMM searches, and phylogenetics to better understand the evidence for lateral gene transfer, gene loss, and rate variation in *ACY3/ASPA* homologues, the most parsimonious explanation for the distribution of *ACY3/ASPA* genes in eukaryotes likely involves both gene loss and lateral gene transfer, albeit lateral gene transfer that occurred hundreds of millions of years ago prior to the divergence of gnathostomes and even longer and prior to the divergence of bilateria. Given the many known, well-characterized, and adaptive lateral gene transfers from bacteria to insects and nematodes, lateral gene transfers at these time scales in the ancestors of humans is expected.

## Background

“If all the trees were one tree, what a great tree that would be.” – Mother Goose

We have one great tree of life that grows and is pruned by evolutionary processes. In 1859, Darwin published “On the Origin of Species” describing the role natural selection plays on the evolution of species, elucidating an interplay between competition and survival [1]. Seventy years later, Frederick Griffith discovered that traits, specifically virulence, can be directly transferred between bacteria in a process we now understand to be horizontal/lateral gene transfer (HGT/LGT) [2]. It was another 16 years before Avery, MacLeod, and McCarty demonstrated that DNA is the molecule that encodes traits and is inherited [3]. Darwin’s theory predates the discovery of DNA and as such transcends any one specific molecular mechanism.

Today, the field of molecular evolution focuses on understanding Darwinian evolution of genomes along with Kimura’s neutral theory with an emphasis on using phylogenetic techniques to analyze nucleotide sequence variation in protein coding genes. With some exceptions, this research in eukaryotes focuses on nucleotide substitutions in conserved protein-coding regions from genes deemed *a priori* to be vertically inherited. But as Avery et al. discovered, traits can also be transferred horizontally or laterally via LGT. LGT has played a major role in the natural evolution and niche adaptation of bacteria, but the role of LGT in the evolution of eukaryotic genomes has been understudied and underappreciated.

When we started working on LGT of bacterial DNA into animal genomes more than a decade ago, the prevailing paradigm was that it was non-existent. Subsequently, instances of bacteria-animal LGT have been observed in multiple invertebrates [4-33], including many such integrations of genes that have at least some evidence for being functional [8-18, 22-25, 27, 29, 31-33]. The coffee berry borer acquired a bacterial mannanase gene that allows it to exploit coffee berries as a new ecological niche relative to its sister taxa [22]. The invasive brown marmorated stink bug that ravages crops in the mid-Atlantic region is thought to have several LGTs from bacteria, including a mannanase gene [13]. Several plant parasitic nematodes have acquired cellulases, pectate lyases, and expansin-like proteins from bacteria that allow them to degrade plant material [25, 27]. In mealybugs, LGTs from at least three different bacterial lineages have resulted in hybrid biosynthetic pathways [12]. There have been numerous functional transfers of bacterial peptidoglycan remodeling genes [10, 11, 13, 31-33] to various eukaryotes that may indicate that eukaryotes can acquire bacterial genes that the eukaryotes then use against the bacteria [34].

Despite this, LGT in multicellular eukaryotes with differentiated tissues, particularly gonads, continues to be met with skepticism. For example, Crisp et al. conducted a detailed examination of 26 animal genomes in order to identify putative LGTs in invertebrate and vertebrate genomes, including the human genome [35]. They found that there are fewer predicted LGTs in vertebrates/chordates than invertebrates, but there is still evidence of LGT into chordates, including humans [35]. Some might not find LGT to chordates to be unusual, since chordates are known to have co-opted endogenous retroviral *env* genes multiple times during the evolution of placental mammals [36]. However, LGT from bacteria is thought to pose a higher barrier than acquisition of new functions from endogenous retroviruses.

More recently, Salzberg re-analyzed a subset of the putative LGTs into vertebrates that were proposed by Crisp et al. [35] and concluded that “horizontal gene transfer is not a hallmark of the human genome” [37]. One of the genes Crisp proposed to be a bacteria-chordate LGT [35], but Salzberg attempts to debunk [37], is an N-acyl-aromatic-L-amino acid amidohydrolase (ENSG00000132744), which encodes ACY3. ACY3 can convert *N*-acyl-aromatic-L-amino acid to the corresponding aromatic-L-amino acid and a carboxylate, or alternatively ACY3 can convert *N*-acetyl-L-cysteine-S-conjugate to L-cysteine-S-conjugate and acetate [38]. ACY3 has an important role in human catalyzing the deacetylation of mercapturic acids in kidney proximal tubules [38]. It is highly expressed in the gastrointestinal tract, the endocervix of women, and the kidneys [39, 40]. BLASTP searches of NR with ACY3 returns matches to the human ASPA protein. ASPA is a protein that converts *N*-acetylaspartate to aspartate and acetate [41]. In humans, mutations in ASPA are responsible for Canavan disease, an autosomal recessive disease leading to brain defects and subsequently early death in children [41]. It is expressed in the central nervous system [39, 40]. Given that ACY3 and ASPA are homologues and our BLASTP searches, and therefore likely the BLASTP searches by Crisp et al. [35] and Salzberg [37], return both homologues, we will refer to them as the ACY3/ASPA homologues.

Salzberg discounted the ACY3/ASPA homologues as “no HGT” [37] because it no longer passes the test Crisp et al. [35] devised for bacteria-chordate LGT since it has a significant BLAST match to the recently deposited genome of *Saccoglossus kowalevskii* [42], a flatworm, Metazoan, and hemichordate. We sought to examine the evolutionary history of the *ACY3/ASPA* homologues further in an effort to better understand the evidence for lateral gene transfer, gene loss, and rate variation.

## Results and Discussion

### Phylogeny of human aspartoacylase

BLASTP was used to identify homologues of the human aspartoacylase gene (ENSG00000132744; NP_542389; ACY3) in the non-redundant protein database (NR) using the NCBI website. This search largely confirmed the BLAST-based results from Crisp [35] and Salzberg [37] demonstrating a large number of matches from bacteria and chordates, but no significant matches (e-value <1e-5) from arthropods, nematodes, plants, fungi, or apicomplexa, among others. This BLAST search identified the proteins encoding human ACY3 and the human ASPA (NP_000040; ASPA), as well as their homologues in other animal genomes. A fast-minimal evolution tree and a neighbor joining tree generated using the NCBI BLAST interface both reveal two major clades, one dominated by proteins from bacteria and one dominated by ASPA and ACY3 proteins from vertebrate animals, including humans. An alignment of these homologous proteins was generated and a limited number of poorly aligned sequences were removed, including truncated isoforms. This final alignment included sequences from hundreds of bacteria and chordates as well as alveolates (n=2), chromophytes (n=6), cnidaria (n=2), and hemichordates (n=1).

A maximum likelihood phylogeny inferred with RAxML after model testing with PROTTEST reveals 88% support for a clade of mostly vertebrate proteins and a clade of mostly bacteria, which initially gives an impression of lateral gene transfer, with a gene moving from bacteria to vertebrates, or vice versa. Further refinement of the tree, however, collapsing branches down to the class reveals that the vast majority of the eukaryotic proteins (**Figure 1**) are evolving in a manner consistent with our understanding of the eukaryote evolution (**Figure 2**). The human ACY3 and ASPA are paralogs that likely arose following duplication. This duplication may have occurred after the divergence of bilateria (88% support) in the ancestor of deuterostomia, which includes chordates, hemichordates, and echinodermata. Alternatively, given the poor support for the position of hemichordate, tunicate, and cephalochordate ACY3/ASPA proteins (<60%), this duplication may have occurred as recently as the ancestor of gnathostomes, which includes the majority of vertebrate animals (**Figure 1**). The latter is consistent with the 1R or the 2R whole genome duplications, which were predicted to occur in the ancestor of hyperoartia(lampreys)/hyperotreti(hagfishes) and the ancestor of gnathostomes, respectively [43]. Alternatively, given the poor support values, ASPA/ACY3 may have been acquired by hemichordates, tunicates, and cephalochordates from another animal early in animal evolution.

**Figure 1.**
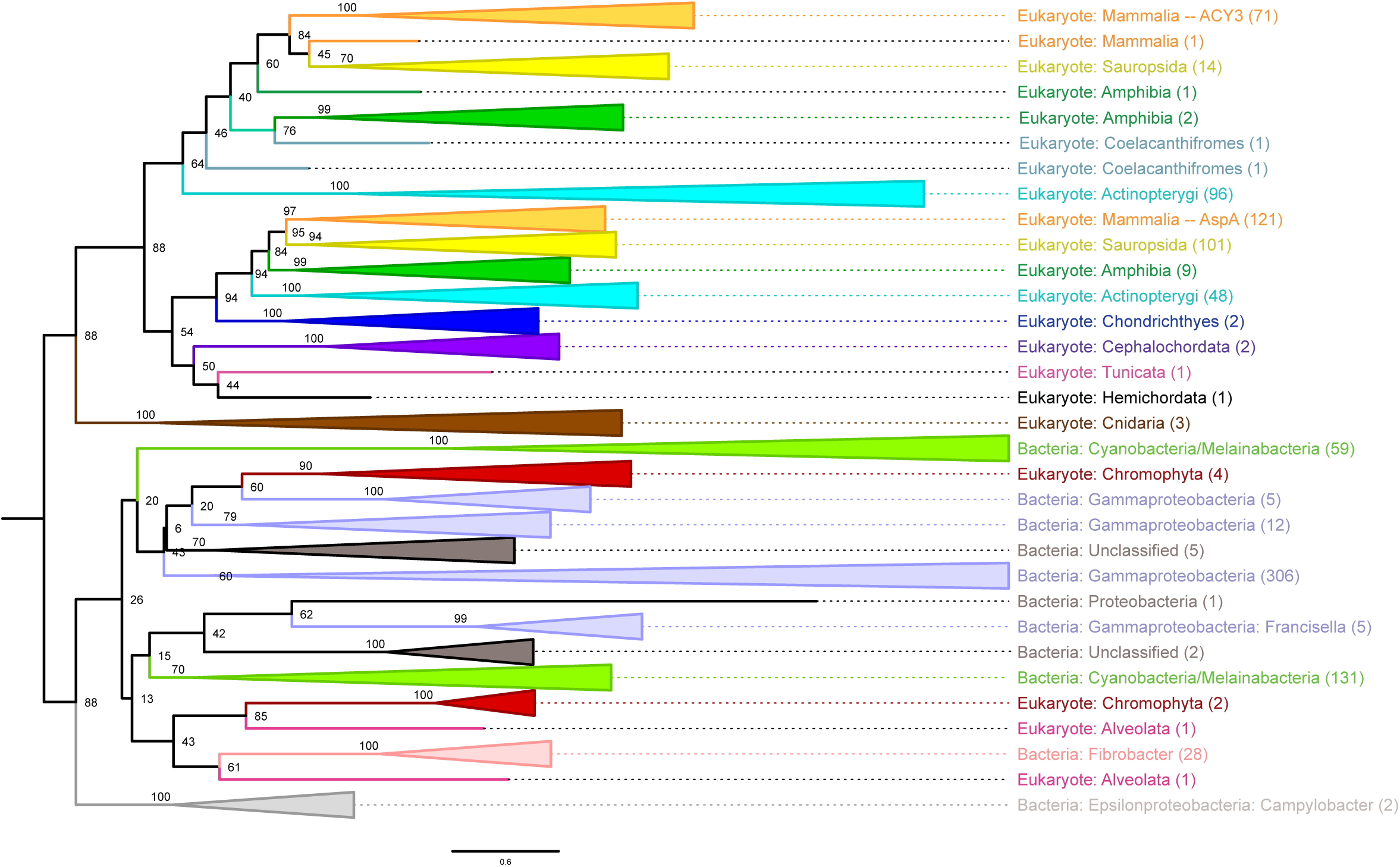
Maximum Likelihood Phylogeny of ACY3/ASPA Homologues. The maximum likelihood (ML) phylogeny of ACY3/ASPA homologues inferred with RAxML is visualized with FigTree in a rectangular phylogram rooted on the edge between the majority of eukaryotic proteins and the majority of prokaryotic proteins. When appropriate and supported by a high support value, branches are collapsed and illustrated with triangles that are color-coded according to the taxonomic distribution of the members. The number of proteins represented in the collapsed branches are noted in parentheses on the right.

**Figure 2.**
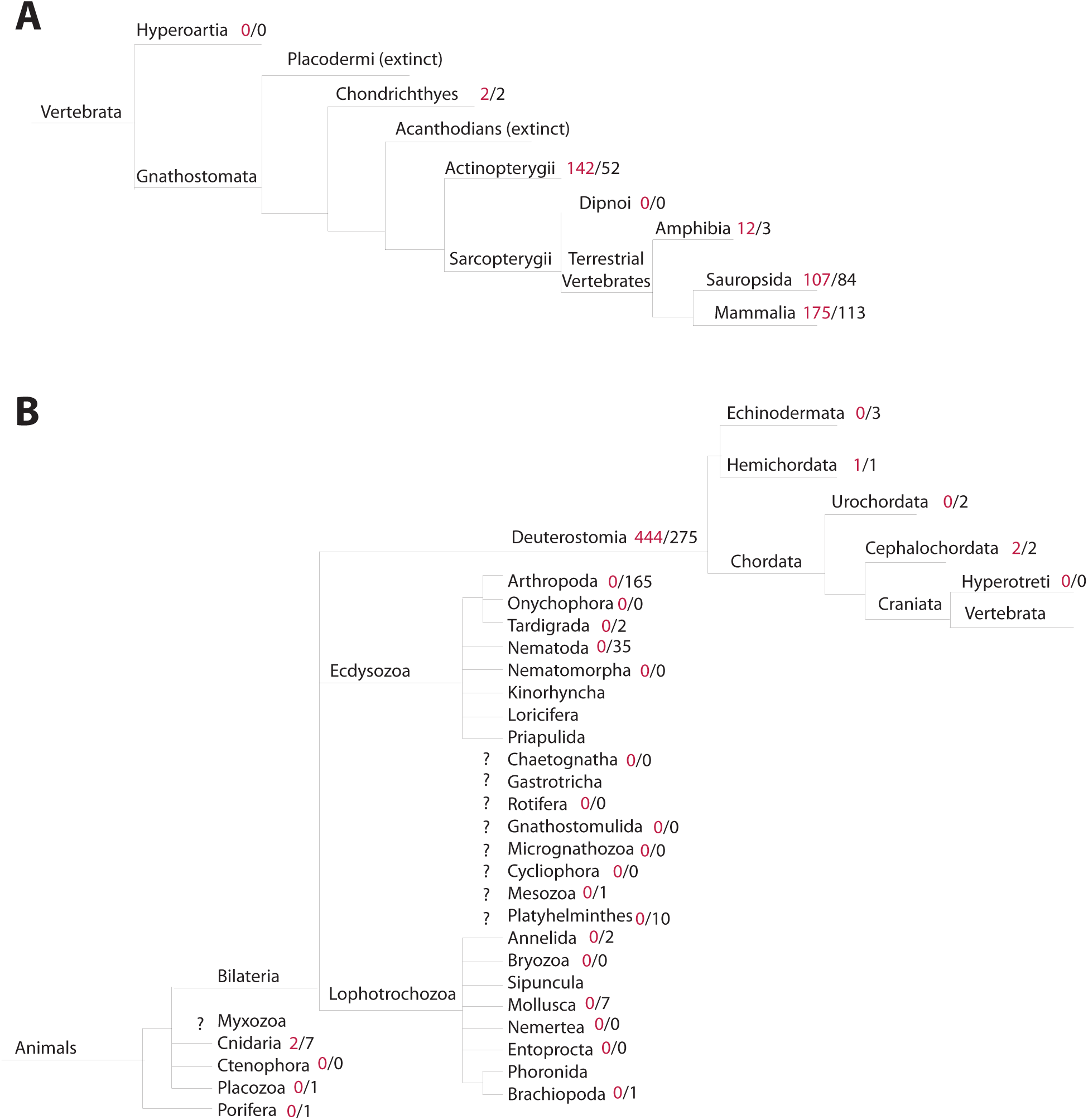
Gene Loss Analysis. Phylogenies from the Tree of Life Web Project were concatenated and used to interpret gene loss. However, it is important to consider that some of the older branches in the tree of life are still disputed. The animal phylogeny was broken out into two panels illustrating: (A) the evolution of mammals from vertebrates and (B) the evolution of vertebrates from animals. To assess gene loss, the number of ACY3/ASPA homologues in a given taxonomic lineage were compared to the number of organisms with >5,000 proteins deposited in public databases. ACY3/ASPA homologues are consistently found in the deuterostome lineage, but are missing from some well-sequenced sister taxa like arthropods and nematodes. There are inadequate levels of genome sequence data at key taxonomic levels to enable the delineation of the relative contribution of LGT, gene loss, and rate variation for ASPA/ACY3 homologues.

The relationship between bacteria and eukaryote proteins is less clear in non-animals. Proteins from chromophytes and alveolates are nested amongst proteins from disparate bacterial taxa (88% support), predominantly cyanobacteria, fibrobacteria, and gamma-proteobacteria, but also two Campylobacter, which are epsilonproteobacteria (**Figure 1**). Chromophytes and alveotates are both Chromista, a group of non-animal/metazoa eukaryotic photosynthetic organisms that likely acquired their chloroplasts from red algae. The structure of the phylogeny suggests that lateral gene transfer may have occurred between the bacterial and these non-animal/metazoan eukaryotic ASPA/ACY3 homologues although the support values for the tree topology do not allow for any further elucidation of the relationship. On one extreme, ACY3/ASPA may have been present in the ancestor of all eukaryotes and was acquired by bacteria via LGT from the ancestor of chromophyta and alveolata, and on the other extermee, ACY3/ASPA may have been acquired by animals, chromophyta, and/or alveolates from bacteria. However, it is important to consider that only extant proteins are examined with a phylogenetic tree, and gene loss is not evaluated.

### Gene loss or lateral gene transfer?

Protein phylogenies only examine the relationship between extant proteins that have been sequenced. Two alternate hypotheses to consider when examining evidence for/against LGT are gene loss and rate variation. To consider gene loss, information about lineages lacking these proteins is required since some taxa are more abundant on earth, and genome sequencing has been unevenly applied across taxa. For example, despite many arthropod and nematode sequences in NR, arthropod and nematode homologues of ACY3/ASPA were not identified in the BLASTP searches. To account for this, the numbers of ACY3/ASPA homologues for given taxonomic levels were compared to the number of organisms at that taxonomic level that have >5,000 protein sequences in NR. If an organism has >5,000 protein sequences in NR, any ACY3/ASPA homologues are likely to have been sequenced and identified through the BLASTP searches of NR. Nearly identical results were obtained for thresholds between 5,000 and 10,000 proteins, giving confidence that a threshold of 5,000 proteins was neither too stringent nor lenient. However, it is important to note that while it is likely that ACY3/ASPA homologues have been sequenced, genome and transcriptome assemblies can be incomplete and as such absence may be overpredicted. Contamination is also a concern, which would lead to underpredicting absence. However, in most cases at least two organisms of a taxa were sequenced and had concordant results.

Among the chordates, ACY3/ASPA homologues are distributed among all of the vertebrate lineages that were sequenced sufficiently (Figure 2A). In all vertebrate lineages, more ACY3/ASPA homologues were identified in NR from that taxon than there were organisms with >5,000 protein sequences in NR for that taxon suggesting that many of the vertebrate organisms contain at least one ACY3/ASPA homologue. This strongly supports the conclusion that the aspartoacylase was likely present in the ancestor of all vertebrates, or at least gnathostomata.

Among the deuterostomes, there is very limited sequencing outside the chordates with only 3 echinodermata, 1 hemichordata, 2 urochordata, 2 cephalochordata, and no hyperotreti with >5,000 proteins characterized in NR (**Figure 2b**). Of those, the hemichordate and the cephalochordate have ACY3/ASPA homologues, and it is the BLAST match to the hemichordate homologue that led to the reassignment of this as “not HGT” by Salzberg. If ACY3/ASPA were present in the ancestor of all deutersomes, then ACY3/ASPA proteins were lost or unsequenced in the 2 urochordata and 3 echinodermata sequenced.

While ACY3/ASPA homologues are prevalent among the deuterostomes, they are noticeably absent in the 202 ecdysozoa with >5,000 proteins in NR, including arthropods, nematodes, and tardigrades. Likewise, they are also absent from the 10 lophotrochozoa with >5,000 proteins in NR, including annelids, brachiopods, and mollusks. They are also missing in unplaced bilateria taxa, including platyhelminths and mesozoa (**Figure 2B**). Therefore, this protein has a limited taxonomic distribution within bilateria such that if the ACY3/ASPA homologue was present in the ancestor of bilateria, it would have needed to be lost from numerous lineages, including at least ecdysozoa, lophotrochozoa, urochordata, and echinodermata, given our current understanding of genomics/transcriptomics and the tree of life.

In animals, there were two ACY3/ASPA homologues not in the bilateria and they were in cnidaria (**Figure 2B**). While there are lots of bilaterian species that have >5,000 proteins in NR, only 9 non-bilateria species have been sequenced across the other five taxa – 7 cnidaria, 1 placozoa, and 1 porifera -- and only 2 cnidaria have an ACY3/ASPA homologue (**Figure 2B**). If the ACY3/ASPA homologue was present in the ancestor of all animals, it would have had to have been lost from some cnidaria as well as placozoa and porifera, in addition to the four bilateria lineages discussed above. Given that extensive lateral gene transfers occur in insect and nematodes, other bilateria, LGT might actually be a more parsimonious explanation than gene loss. In other words, two LGTs of a bacterial gene into animals, one in cnidaria and one in deuterostomes, could be more likely than a half dozen gene loss events across diverse animal taxa. Unfortunately without rate estimates for both gene loss and LGT it is hard to be definitive.

### Rate variation

Another alternate explanation to lateral gene transfer and/or gene loss is rate variation. When considering rate variation, some proteins are under accelerated rates of evolution relative to other proteins. For example, following duplication two proteins may diverge at different rates. Following LGT, genes might be expected to undergo different rates of evolution as they enter a new environment. As such rate variation and LGT are not mutually exclusive. However, when considering rate variation as an alternative to LGT we are looking for signatures that might suggest that one lineage has vertically inherited genes that are evolving at a different rate confounding BLAST- and phylogeny-based methods.

To examine this, we relied on the results of two large pre-computed datasets, eggNOG and PFAM. EggNOG is an algorithm that currently uses graph-based unsupervised clustering to identify orthologous genes in 2,031 eukaryote and prokaryote genomes. When eggNOG is interrogated with ACY3 or ASPA, an orthologous group is identified that is found to contain proteins from bacteria and metazoans, making it largely similar to the results returned with the BLAST-based results described above. However, key taxa were not recovered like the hemichordate homologues, likely because it was only recently reported.

PFAM uses hidden Markov model (HMM) searches to find functionally-related, but substantially-diverged, proteins. HMM searches rely on the use of probabilistic, hidden Markov models to identify protein homologues with great sensitivity and specificity; these models quickly and efficiently find homologues based on the presence of protein features shared between homologues (e.g. catalytic residues) not identified through traditional BLAST-based searches. These HMM results can be overlaid on a species tree (e.g. for Acy3/ASPA: http://pfam.xfam.org/family/AstE_AspA#tabview=tab7).

Among the metazoa, the HMM searches yielded the same results as the gene loss analysis above, except for a match to *Acyrthosiphon pisum* (pea aphid) which is an arthropod. The match is not to a protein in NR, but instead to a nearly 1 kbp region on a 1.2 kbp contig from the whole genome sequencing project that has closest similarity to proteins annotated as succinylglutamate desuccinylase from *Pantoea* endosymbionts. It is likely that this is a contig from a contaminant bacterial endosymbiont given that it matches at 92% nucleotide identity to the complete genome of *Pantoea rwandensis* strain ND04 over 100% of the length of the 1.2 kbp *A. pisum* contig.

However, in more distant eukaryotic lineages, the HMM gave different results from eggNOG and BLAST. More than 3,000 sequences from ACY3/ASPA homologues can be found in 1,583 species including many eukaryotes as well as archaea suggesting a broad taxonomic distribution of organisms with this domain. Two taxonomically disparate plant taxa, *Cajanus cajan* (pigeon pea) and *Monoraphidium neglectum* (single-cell green alga) as well as 49 fungi across many diverse fungal lineages contain proteins with the ACY3/ASPA domain. Unfortunately, and similar to the problem with phylogenetic trees, no information can be gleaned about taxa lacking ACY3/ASPA domain-containing homologues. However, it is clear that the functional domain exists in taxa beyond those identified with BLAST or eggNOG searches suggesting that there can be substantial sequence divergence. However, the sequence divergence and our inability to produce high quality alignments of these sequences precludes further analyses beyond this limited analysis of their taxonomic distribution.

## Conclusions

Salzburg [37] and others have stated, when referring to LGT, that extraordinary claims require extraordinary evidence, implying that LGT is an extraordinary claim. Salzberg goes on to suggest that more mundane explanations are at play, like gene loss and rate variation [37]. It seems unlikely that gene loss and rate variation alone can explain these results. On one extreme, it seems reasonable that eukaryotes may have acquired these genes from bacteria a handful of times, once in the ancestor of fungi and at least once in animals as well as an unresolved number of times in alveolates/chromophytes. On the other extreme, it also seems equally reasonable that instead, the genes have been vertically inherited in eukaryotes with dozens of gene loss events and at least one lateral gene transfer to bacteria, where it could have spread further via lateral gene transfer. It is not possible to be more definitive at this time given the lack of sufficient genome sequencing of key taxa, like hyperartia, hyperotreti, ctenophore as well as more sequence data from placozoa, porifera, cephalochordate, urochordata, hemichordate, and echniodermata. However, the most parsimonious explanation for the distribution of ACY3/ASPA homologues involves some bacteria-animal lateral gene transfer. This case, of a gene essential for proper brain development and function that seems to have a limited phylogenetic distribution, illustrates how these comparisons need careful scrutiny. It highlights the need for more robust, focused analyses on the extent of LGT, gene loss, and rate variation in eukaryotes and their influence on trait acquisition. Furthermore, unbiased estimates of LGT and gene loss rates across and between different taxa are desperately needed to understand the likelihood of both events. Lastly, our understanding of the topology of the tree of life also influences these analyses, and many important branches have yet to be resolved or remain in dispute. Collectively, this analysis demonstrates our need for further high quality complete genome and transcriptome assemblies from key phylogenetic groups in order to have the power to infer the correct relationships between both taxa and proteins of interest in order to properly evaluate claims of LGT and gene loss.

## Methods

### BLAST searches

ACY3/ASPA homologues were identified from a BLASTP search [44] using ACY3 as a query (ENSG00000132744; NP_542389; ACY3) and NR as a reference using the NCBI BLAST server during August and September 2017. A similar search using ASPA as the query produces similar results, but all subsequent analyses were conducted on the output using ACY3 as a query. All BLASTP searches were performed with the default parameters except that 20,000 results were allowed to be returned. To identify homologues in specific clades, the BLASTP searches were restricted to these clades using the appropriate taxon_id (e.g. fungi/taxid:4751, plants/taxid:3193, arthropods/taxid:6656, insects/taxid:6960, nematodes/taxid:6231, mollusks/taxid:6447, and apicomplexan/taxid:5794). A neighbor joining tree and a fast-minimum evolution tree were generated using the NCBI BLAST interface with maximum sequence difference of 0.85 and Grishin distance labeling sequences by taxonomic name.

### Multiple sequence alignment, model testing, and inferring/visualizing phylogenetic trees

All of the protein sequences identified from the ACY3-based BLASTP searches were downloaded locally and aligned with CLUSTALW v.1.4 [45] as implemented in Bioedit v.7.2.5 [46]. Poorly aligned sequences, particularly partial sequences and isoforms, were removed manually. The sequences were then re-aligned with CLUSTALW v.1.4 [45] as implemented in Bioedit v.7.2.5 [46]. The best-fit model of amino acid substitution was determined for each of the datasets with ProtTest3.2 [47]. All 15 models of protein evolution were tested in addition to the +G parameter (i.e. including models with rate variation among sites). RAxML v.8.2.10 [48] automatically removed undetermined columns and sequence duplicates and was used to infer the phylogeny with 1000 rapid bootstrap inferences, a thorough ML search, the GAMMA model of rate heterogeneity, the ML estimate of alpha-paramter, and the JTT substitution matrix using the command raxmlHPC -f a -m PROTGAMMAJTT -p 12345 -x 12345 -N autoMRE. Taxonomic information from the NCBI Taxonomy database was added to the RAxML output. Accessions that lack an entry in the taxonomy database were left blank. However, in some figures the genus and species designations were added in manually after confirming the lack of an entry in the taxonomy database; these are denoted with an asterisk (*). Phylogenetic trees were visualized with Dendroscope v.3.5.7 [49].

### Gene Loss

Taxa with >5,000, >6,000, >7,000, >8,000, and >10,000 known proteins in NR were determined by using the NCBI protein server (https://www.ncbi.nlm.nih.gov/protein) to search PDB, RefSeq, UniProtKB/Swiss-Prot, DDBJ, EMBL, GenBank, and PIR with the appropriate taxon_id in November 2017. These results were overlaid on reference phylogenetic trees for the eukaryotic lineages that were concatenated from trees retrieved from the Tree of Life website (tolweb.org) [50-61].

### Rate variation

In order to identify ACY3/ASPA homologues that may be subject to rate variation, pre-computed orthologous clusters in eggNOG were examined (http://eggnogdb.embl.de/#/app/results#COG2988%5Fdatamenu) as well as hidden markov model (HMM) search results generated by PFAM were overlaid on a species tree using the PFAM server (http://pfam.xfam.org/family/AstE_AspA#tabview=tab7).

## List of Abbreviations

HGT: horizontal gene transfer
HMM: hidden Markov model
LGT: lateral gene transfer
ML: maximum likelihood

## Declarations

### Ethics approval and consent to participate

Not applicable.

### Consent for publication

Not applicable.

### Availability of data and materials

The phylogenetic tree generated in this analysis will be deposited in Dryad with an accession and permanent identifier. However, Dryad does not allow for submission of materials until a manuscript is accepted for publication.

### Author contributions

JCDH conducted all analyses and wrote the manuscript.

## Acknowledgements

We would like to thank John Werren at the University of Rochester for helpful discussions, James Munro at the Institute for Genome Science for his help with running PROTTEST, and John Mattick for his help adding taxonomy information to the phylogenetic trees from RAxML at the Institute for Genome Science. This work was funded by the National Science Foundation Advances in Biological Informatics (ABI-1457957) and an NIH Director’s Transformative Research Award (1-R01-CA206188). to JCDH.

